# Time-dependent enhancement of mRNA vaccines by 4-1BB costimulation

**DOI:** 10.1101/2024.03.01.582992

**Authors:** Sarah Sanchez, Tanushree Dangi, Bakare Awakoaiye, Nahid Irani, Slim Fourati, Justin Richner, Pablo Penaloza-MacMaster

## Abstract

mRNA vaccines have demonstrated efficacy against COVID-19. However, concerns regarding waning immunity and breakthrough infections have motivated the development of next-generation vaccines with enhanced efficacy. In this study, we investigated the impact of 4-1BB costimulation on immune responses elicited by mRNA vaccines in mice. We first vaccinated mice with an mRNA vaccine encoding the SARS-CoV-2 spike antigen like the Moderna and Pfizer-BioNTech vaccines, followed by administration of 4-1BB costimulatory antibodies at various times post-vaccination. Administering 4-1BB costimulatory antibodies during the priming phase did not enhance immune responses. However, administering 4-1BB costimulatory antibodies after 96 hours elicited a significant improvement in CD8 T cell responses, leading to enhanced protection against breakthrough infections. A similar improvement in immune responses was observed with multiple mRNA vaccines, including vaccines against common cold coronavirus, human immunodeficiency virus (HIV), and arenavirus. These findings demonstrate a time-dependent effect by 4-1BB costimulation and provide insights for developing improved mRNA vaccines.

## INTRODUCTION

mRNA vaccines prevent severe SARS-CoV-2 infection and are being explored for multiple diseases such as influenza, HIV, and cancer ^1–3^. While mRNA vaccines have shown high efficacy at preventing COVID-19, they do not always confer sterilizing immunity against symptomatic SARS-CoV-2 infection. Concerns also linger about the durability of immune responses after mRNA vaccination, and the high prevalence of breakthrough infections among vaccinated individuals underscores the need for improved mRNA vaccine formulations.

T cells are critical for controlling early viral dissemination and their function is associated with reduced disease severity following breakthrough SARS-CoV-2 infection ^4–6^. The activation of T cells relies on two crucial signals: First, naïve T cells require specific antigen recognition via their T cell receptor (TCR), and second, they require costimulation. The most widely studied costimulatory pathway is CD28/B7, but the effects of other less-known costimulatory pathways like 4-1BB/4-1BBL remain poorly understood. 4-1BB (also known as CD137) is a costimulatory receptor that belongs to the tumor necrosis factor receptor (TNFR) superfamily and is important for T cell responses. 4-1BB is expressed mostly on T cells and NK cells, among other cells, whereas its ligand (4-1BBL) is expressed mostly on antigen presenting cells ^7, 8^. 4-1BB costimulation is important for T cell responses following bacterial and viral infections, and triggering of this pathway results in increased T cell proliferation, survival, and effector functions ^9–13^. Prior studies have also shown that 4-1BB costimulation renders effector T cells resistant to suppression by T regulatory cells ^14^, and due to its immunostimulatory effects, 4-1BB costimulatory antibodies have been explored in cancer immunotherapy in several human trials ^15–18^.

Various studies have demonstrated an immunostimulatory role for 4-1BB, but there is also evidence for an immunoinhibitory role. For example, 4-1BB-/- mice exhibit exaggerated primary CD8 T cell responses following cytomegalovirus (CMV) infection, and administration of 4-1BB costimulatory antibodies before viral infection results in impaired CD8 T cell responses ^19, 20^. 4-1BB costimulatory antibodies can also abrogate autoimmunity during autoimmunity ^21, 22^. Thus, 4-1BB costimulation can play context-dependent roles on immune responses following viral infection, cancer progression, and autoimmunity, sometimes stimulating immune responses, and sometimes suppressing immune responses. The effects of 4-1BB costimulation on vaccine responses, however, remain understudied. Here, we studied the time-dependent effects of 4-1BB costimulation on immune responses elicited by mRNA vaccines. We show that triggering 4-1BB costimulation at day 4 post-vaccination significantly improves the efficacy of mRNA vaccines, rendering these vaccines better able to prevent subsequent breakthrough infections. These studies highlight a novel strategy to improve mRNA vaccines via time-dependent modulation of the 4-1BB pathway.

## RESULTS

### Time-dependent effects of 4-1BB costimulation on CD8 T cell responses

In this study, we attempted to improve mRNA vaccines by triggering a T cell costimulatory pathway, known as 4-1BB. First, we conducted experiments to investigate how the timing of 4-1BB costimulation affects vaccine efficacy following mRNA vaccination. We first primed C57BL/6 mice intramuscularly with an mRNA lipid nanoparticle vaccine expressing the SARS-CoV-2 spike protein (mRNA-spike), and on the same day we injected these mice with 4-1BB costimulatory antibodies or control antibodies to examine the effect of triggering 4-1BB costimulation during the priming phase (**Fig. S1A**). Treatment of mice with 4-1BB costimulatory antibodies during the priming phase (day 0 of vaccination) did not improve CD8 T cell responses, relative to control (**Fig. S1B**), demonstrating that simultaneous antigenic recognition (Signal 1) and 4-1BB costimulation (Signal 2) on the same day did not improve CD8 T cell responses. In fact, there was a severe impairment in antibody responses in mice that received 4-1BB costimulatory antibodies on the same day of vaccination (**Fig. S1C**). These initial data suggested that 4-1BB costimulation at the time of immune priming was detrimental.

As mentioned earlier, antigen recognition and costimulation are signals necessary to generate T cell responses, but the optimal timing between these two signals remains unknown. To answer this simple question, we administered 4-1BB costimulatory antibodies during the later effector phase (day 4) of the immune response (**Fig. 1A**). Interestingly, treatment of mice with 4-1BB costimulatory antibodies at day 4 post-vaccination resulted in a potent increase of vaccine-elicited CD8 T cell responses in blood (**Fig. 1B-1C**) and tissues (**Fig. 1D-1F**). This potentiation of CD8 T cell responses was associated with higher Ki67 expression, suggesting enhanced proliferation (**Fig. 2A-2B**). In addition, treatment with 4-1BB costimulatory antibodies at day 4 post-vaccination resulted in a significant increase in cytokine expression (**Fig. 2C-2D**), and an increase in systemic cytokines, including IFNγ, MIP-1α, GM-CSF, and TNFα relative to control vaccination (**Fig. 2E-2G**).

**Fig. 1.**
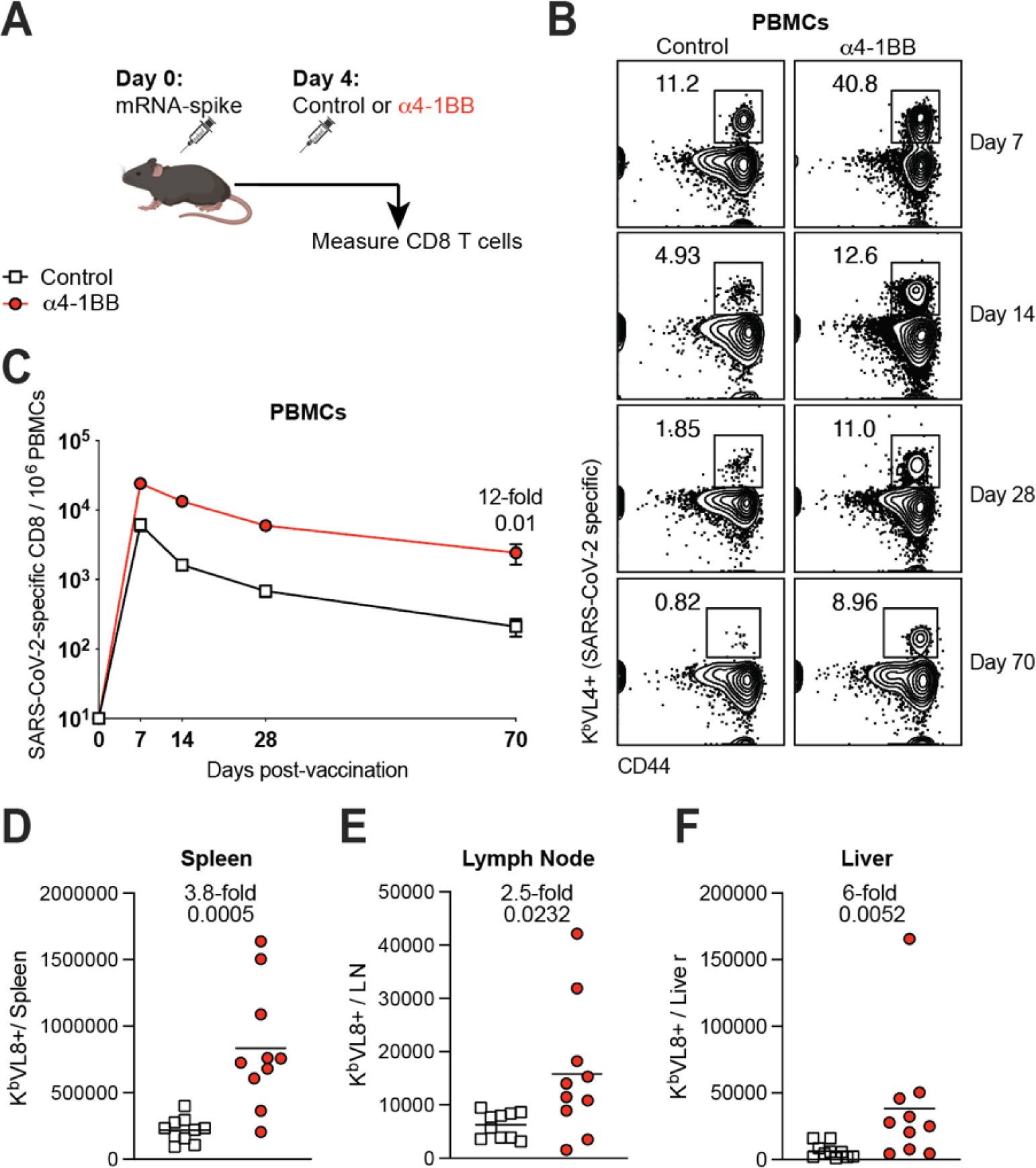
4-1BB costimulation during the early effector phase improves CD8 T cell responses following mRNA-SARS-CoV-2 vaccination. **(A)** Experimental outline for evaluating whether treatment with α4-1BB improves immune responses elicited by an mRNA-SARS-CoV-2 vaccine in C57BL/6 mice. Mice were immunized with 3 μg of a SARS-CoV-2 mRNA spike vaccine followed by treatment with 50 μg of α4-1BB or control antibodies at day 4. **(B)** Representative FACS plots showing the frequencies of SARS-CoV-2-specific CD8 T cells (K^b^VL8+) in PBMCs. **(C)** Summary of SARS-CoV-2-specific CD8 T cell responses in PBMCs. **(D)** Summary of SARS-CoV-2-specific CD8 T cell responses in spleen. **(E)** Summary of SARS-CoV-2-specific CD8 T cell responses in draining lymph nodes. **(F)** Summary of SARS-CoV-2-specific CD8 T cell responses in liver. Data are from two experiments, n=5 per group/experiment. All data are shown. Indicated *P* values were determined by parametric test (unpaired t test).

**Fig. 2.**
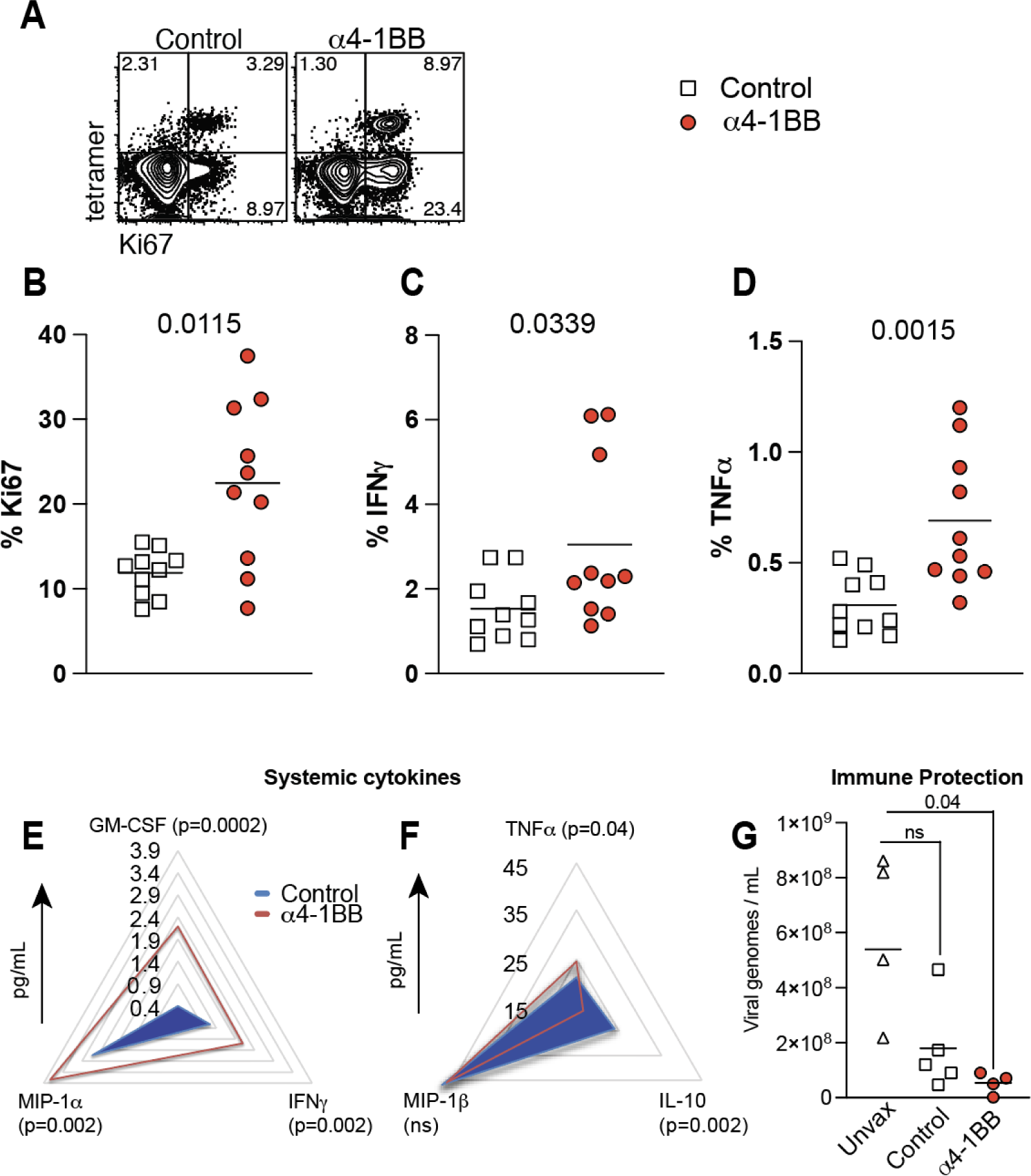
4-1BB costimulation during the early effector phase improves CD8 T cell proliferation and function. Experimental outline was similar to Fig. 1. (**A)** Representative FACS plots showing the frequencies of Ki67 expressing SARS-CoV-2 specific CD8 T cells. **(B)** Summary of SARS-CoV-2-specific CD8 T cells that express the proliferation marker Ki67. In panels C-D, splenocytes were incubated with a SARS-CoV-2 spike peptide (VNFNFNGL) for 5 hr at 37°C in the presence of GolgiStop and GolgiPlug to detect cytokine-expressing CD8 T cell responses. **(C)** Summary of SARS-CoV-2-specific CD8 T cells that express IFNγ. **(D)** Summary of SARS-CoV-2-specific CD8 T cells that express TNFα. **(E-F)** Radar plots showing cytokines in serum 6 hr after treatment with 4-1BB costimulatory antibodies (4-1BB costimulatory antibodies were administered at day 4 post-vaccination, same as in Fig. 1). (**G**) Viral loads in lungs on day 3 post-challenge. K18-hACE2 mice were immunized with 3 μg of a SARS-CoV-2 mRNA spike vaccine followed by α4-1BB or control antibodies at day 4. Mice were challenged intranasally with 5 × 10^4^ PFU of SARS-CoV-2 after 14 days. Viral RNA was quantified by RT-qPCR in lungs. Challenges were performed with a total of 4-5 mice per group in BSL-3 facilities. All other data are from two experiments, n=5 per group/experiment. All data are shown. Indicated *P* values were determined by parametric test (unpaired t test), except for panel G, which used 2-way ANOVA (Dunnett’s multiple comparisons tests).

To measure vaccine protection, we utilized K18-hACE2 mice, which are highly susceptible to SARS-CoV-2. These mice were first vaccinated with the mRNA-spike vaccine and then treated with 4-1BB costimulatory antibodies or control antibodies at day 4. After 2 weeks, these mice were challenged intranasally with SARS-CoV-2, and breakthrough viral loads were measured in lungs at day 3 post-challenge. This challenge study was focused on evaluating viral control at a very early time after infection. Note that all vaccinated mice completely clear infection by day 7 post-challenge, so differences in viral control could only be observed at very early times post-challenge ^4, 23–25^. Importantly, mice that received 4-1BB costimulation at day 4 post-vaccination exhibited a significant reduction in viral loads after SARS-CoV-2 challenge, whereas control vaccinated mice did not show a statistically significant reduction in viral loads, relative to unvaccinated mice. These data suggested that 4-1BB costimulation could improve the efficacy of mRNA-SARS-2 vaccines, allowing the vaccine to protect better against breakthrough infection.

In all the studies above, we administered a single low dose of 4-1BB costimulatory antibodies of 50 μg, like in our prior publication ^10^. Higher and repetitive doses of 4-1BB costimulatory antibodies (on days 4, 7, 10) did not further improve CD8 T cell responses (**Fig. S2A-S2B**). Administration of 4-1BB costimulatory antibodies at day 4 (either at the standard low dose or the high dose) did not significantly affect antibody responses (**Fig. S2C**). These results demonstrate that reinforcement of 4-1BB costimulation, exactly at day 4 post-vaccination, improves CD8 T cell responses.

### Effects of 4-1BB costimulation on CD8 T cell differentiation

Following an initial antigen encounter, CD8 T cells differentiate into distinct subsets, including effector cells (Teff), effector memory cells (Tem), and central memory cells (Tcm) ^26–28^. To evaluate whether 4-1BB costimulation biased the differentiation of specific cell subsets, we FACS-sorted splenic SARS-CoV-2-specific CD8 T cells at day 7 post-vaccination and performed RNA-sequencing (RNA-seq). Interestingly, by principal component analyses, SARS-CoV-2 specific CD8 T cells clustered differently in treated mice, suggesting transcriptional differences (**Fig. 3A**). We observed enrichment of genes associated with cell proliferation, activation, and effector differentiation (**Fig.3B-3E**), and more pronounced effector signatures by gene set enrichment analyses (GSEA) (**Fig. 3F**) in mice that received 4-1BB costimulatory antibodies.

**Fig. 3.**
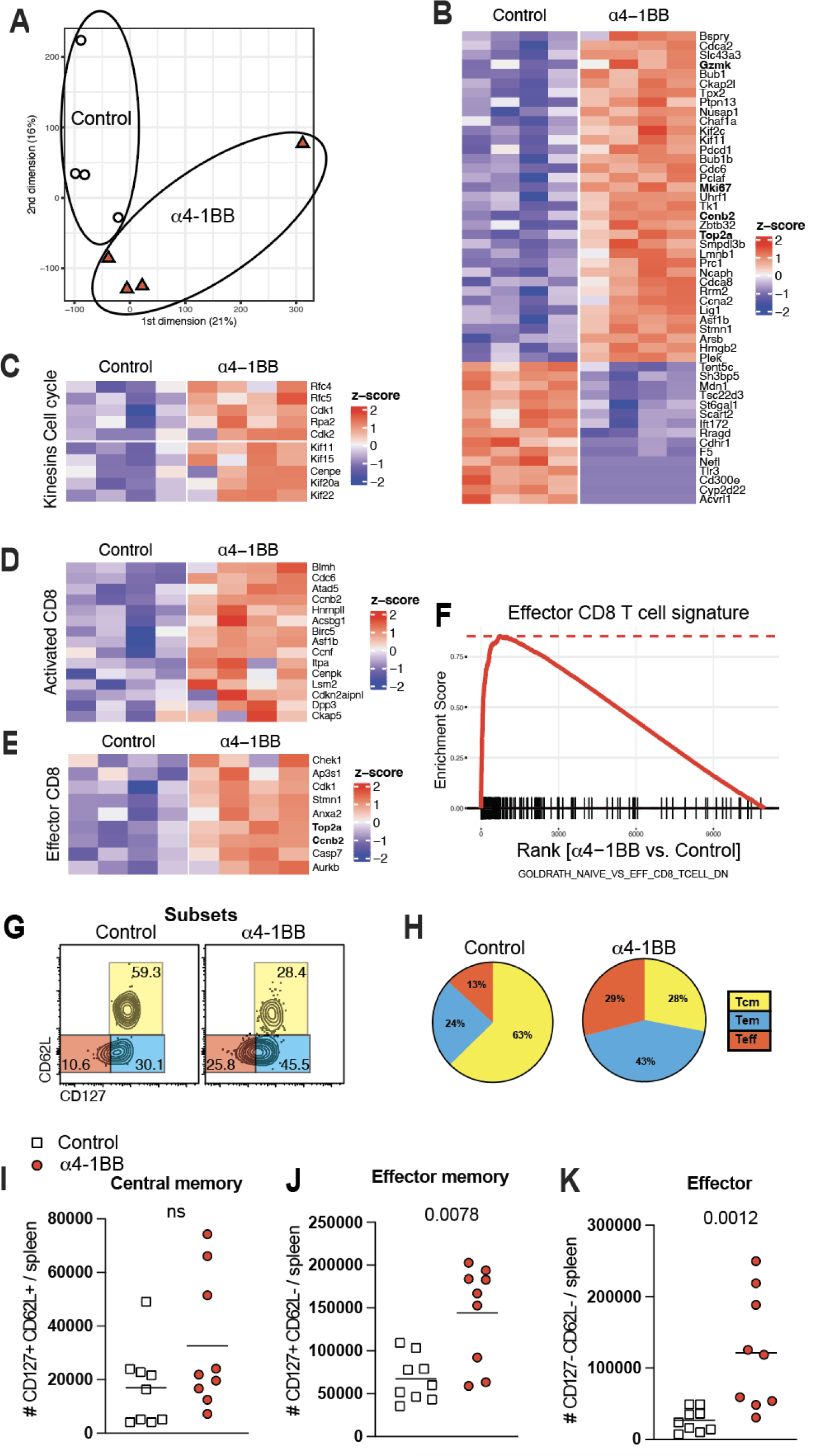
CD8 T cell subset differentiation following 4-1BB costimulation. Experimental outline was similar to Fig. 1. At day 7, splenic CD8 T cells were MACS sorted. Subsequently, live, CD8+, CD44+, K^b^VL8+ cells were FACS-sorted to ∼99% purity and used for bulk RNA-seq. **(A)** Dimension reduction plots showing transcriptional differences. **(B)** Heatmap showing row-standardized expression of selected proliferation (top) and apoptotic genes. **(C)** Heatmap showing row-standardized expression of selected cell cycle (top row) and kinesins (bottom row) genes. **(D)** Heatmap showing row-standardized expression of selected activation genes. **(E)** Heatmap showing row-standardized expression of selected effector genes. **(F)** GSEA plots showing enrichment of effector genes. (**G**) Representative FACS plots showing the frequencies of SARS-CoV-2-specific CD8 T cells (K^b^VL8) that differentiate into effector, effector memory, and central memory T cell subsets. (**H**) Pie diagrams showing the frequencies of CD8 T cell subsets. (**I-K**) Numbers of central memory, effector memory, and effector CD8 T cells. All data are from tetramer+ (K^b^VL8+) cells from spleen. RNA-Seq data are from one experiment with n=4 per group. All other data are from two experiments with n=4-5 per group/experiment. All data are shown. Indicated *P* values were determined by parametric test (unpaired t test).

We validated these gene expression profiling results by performing phenotypic characterization of SARS-CoV-2-specific CD8 T cells using multiparameter flow cytometry. Consistent with the gene expression profiling, CD8 T cells after 4-1BB costimulation exhibited more pronounced effector (CD62L-CD127-) and effector memory (CD62L-CD127+) differentiation (**Fig. 3G-3H**). There were no differences in the total number of central memory CD8 T cells (CD62L+ CD127+) (**Fig. 3I**), but the total number of effector CD8 T cells and effector memory CD8 T cells was greater in mice that received 4-1BB costimulatory antibodies (**Fig. 3J-3K**). These data suggested that 4-1BB costimulation at day 4 selectively expands effector and effector memory CD8 T cells.

### Effect of 4-1BB costimulation on other vaccines

We then interrogated the generalizability of our observations using other mRNA-based vaccines. We recently developed a novel mRNA-based vaccine expressing the spike protein of OC43, a common cold coronavirus that causes recurrent infections in humans. We tested whether administration of 4-1BB costimulatory antibodies at day 4 improves the efficacy of this mRNA-based coronavirus vaccine. Consistent with the prior data, 4-1BB costimulation at day 4 post-vaccination also resulted in a significant improvement in OC43-specific CD8 T cell responses, and no significant differences in antibody responses (**Fig. 4A-4C**). Similar effects were observed with other mRNA-based vaccines, one against HIV (**Fig. 4D-4F**) and another one against ovalbumin (OVA) (**Fig. 4G-4I**). To evaluate immune protection, we challenged mRNA-OVA immunized mice intravenously with a supralethal dose of listeria monocytogenes expressing OVA followed by quantification of bacterial titers in spleen at day 3 post-challenge (**Fig. S3A**). Importantly, mice that received 4-1BB costimulation after vaccination showed sterilizing immunity following challenge, whereas control vaccinated mice showed high bacterial loads (**Fig. S3B**).

**Fig. 4.**
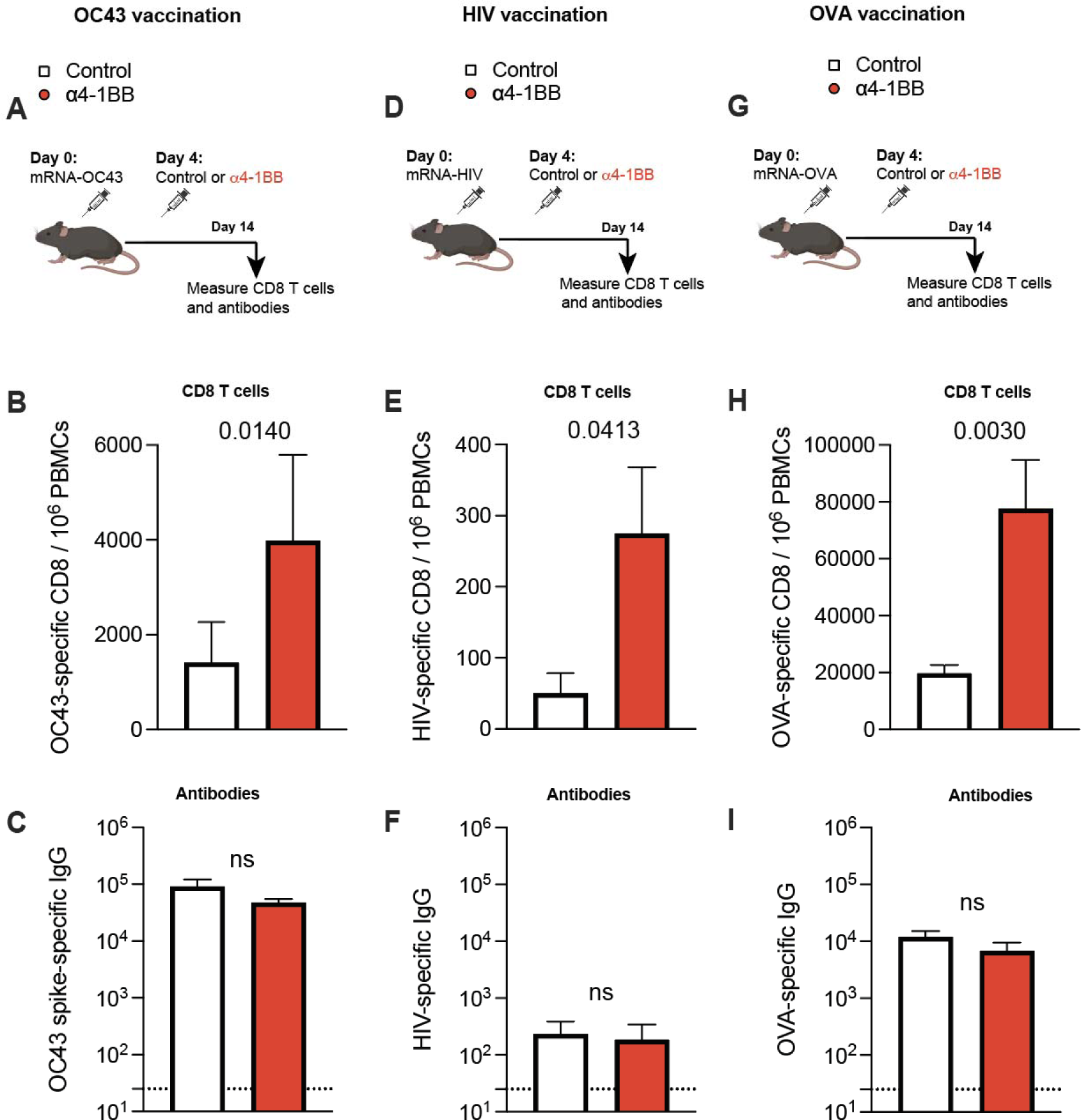
Generalizability to other mRNA vaccines: common cold coronavirus, HIV, and OVA. **(A)** Experimental outline for evaluating whether treatment with α4-1BB improves immune responses elicited by an mRNA-OC43 spike vaccine. **(B)** Summary of OC43-specific CD8 T cell responses in PBMCs. **(C)** Summary of OC43-specific antibody responses in sera. **(D)** Experimental outline for evaluating whether treatment with α4-1BB improves immune responses elicited by an mRNA-HIV (SF162) envelope vaccine. **(E)** Summary of HIV-specific CD8 T cell responses in PBMCs. **(F)** Summary of HIV-specific antibody responses in sera. **(G)** Experimental outline for evaluating whether treatment with α4-1BB improves immune responses elicited by an mRNA-OVA vaccine. **(H)** Summary of OVA-specific CD8 T cell responses in PBMCs. **(I)** Summary of OVA-specific antibody responses in sera. Mice were immunized with 3 μg of each respective vaccine followed by treatment with 50 μg of α4-1BB or control antibodies at day 4. Data are from two experiments, one with n=5 per group/experiment and another one with n=2-3 per group/experiment. All data are shown. Indicated *P* values were determined by parametric test (unpaired t test).

We recently developed an mRNA vaccine based on the lymphocytic choriomeningitis virus (LCMV) glycoprotein (mRNA-GP). Similar to the data above, 4-1BB costimulation at day 4 post-vaccination resulted in a significant improvement in virus-specific CD8 T cell responses specific for the GP33 and GP276 epitopes (**Fig. 5A-5D**). To measure immune protection, we challenged these mice intravenously with chronic LCMV Cl-13 and then quantified viral loads in blood and spleen. Mice that received 4-1BB costimulation at day 4 post-vaccination did not show weight loss after LCMV Cl-13 challenge (**Fig. 5E**), and most of these mice exhibited sterilizing immune protection (**Fig. 5E-5G**). Altogether, our data with multiple mRNA vaccines show that triggering 4-1BB costimulation at day 4 post-vaccination significantly increases vaccine efficacy.

**Fig. 5.**
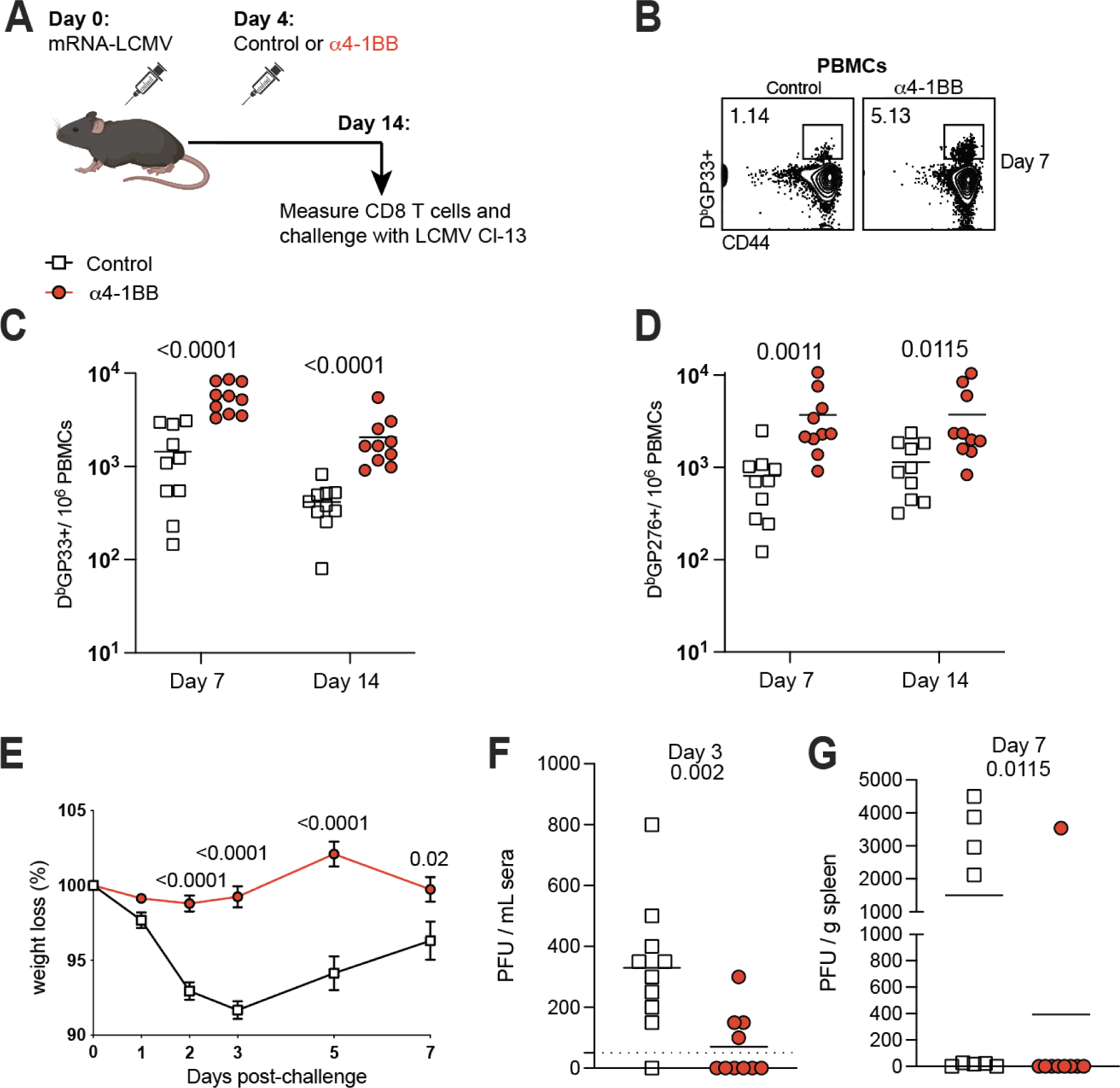
4-1BB costimulation following immunization with an mRNA-based arenavirus vaccine confers sterilizing immunity in most animals. **(A)** Experimental outline for evaluating whether treatment with α4-1BB improves immune responses elicited by an mRNA-LCMV GP vaccine. Mice were vaccinated with 3 μg of an mRNA-LCMV GP vaccine followed by treatment with 50 μg of α4-1BB or control antibodies at day 4. **(B)** Representative FACS plots showing the frequencies of LCMV-specific CD8 T cells (D^b^GP33) in PBMCs at day 7. **(C)** Summary of GP33-specific CD8 T cell responses in PBMCs. **(D)** Summary of GP276-specific CD8 T cell responses in PBMCs. **(E)** Summary of weight loss following chronic LCMV Cl-13 challenge. Mice that received 4-1BB costimulatory antibodies after vaccination were better protected upon chronic LCMV Cl-13 challenge and exhibited no weight loss. **(F)** Viral loads in sera on day 3 post-challenge (after day 7 post-challenge, viral loads were undetectable in the sera in all vaccinated mice). **(G)** Viral loads in spleen on day 7 after infection. Mice were challenged intravenously with 2 × 10^6^ PFU of LCMV Cl-13, and viral loads were quantified by plaque assay. Data are from two experiments, with n=4-5 per group/experiment. All data are shown. Indicated *P* values were determined by parametric test (unpaired t test).

As of now, all our studies have been with mRNA vaccines. We then interrogated whether the effects of “delayed 4-1BB costimulation” could be recapitulated with a different vaccine platform. To answer this simple question, we utilized a poxvirus vector used in the clinically approved Mpox vaccine, based on modified vaccinia Ankara (MVA). Consistent with our prior studies with mRNA vaccines, we also observed improvement of poxvirus-specific CD8 T cell responses when mice were treated with 4-1BB costimulatory antibodies at day 4 post-vaccination (**Fig. 6A-6B**). No difference was observed in poxvirus-specific antibody responses (**Fig. 6C**). Lastly, we extended our results to an MVA-vectored vaccine expressing the SARS-CoV-1 spike antigen derived from the original coronavirus of 2004 (MVA-SARS-1 spike) (**Fig. 6D**). In this model, we also observed improvement of virus-specific CD8 T cell responses when mice were treated with 4-1BB costimulatory antibodies at day 4 post-vaccination (**Fig. S6E-S6F**). Taken together, we show in multiple vaccine models that 4-1BB costimulation at day 4 post-vaccination results in a significant improvement of CD8 T cell responses.

**Fig. 6.**
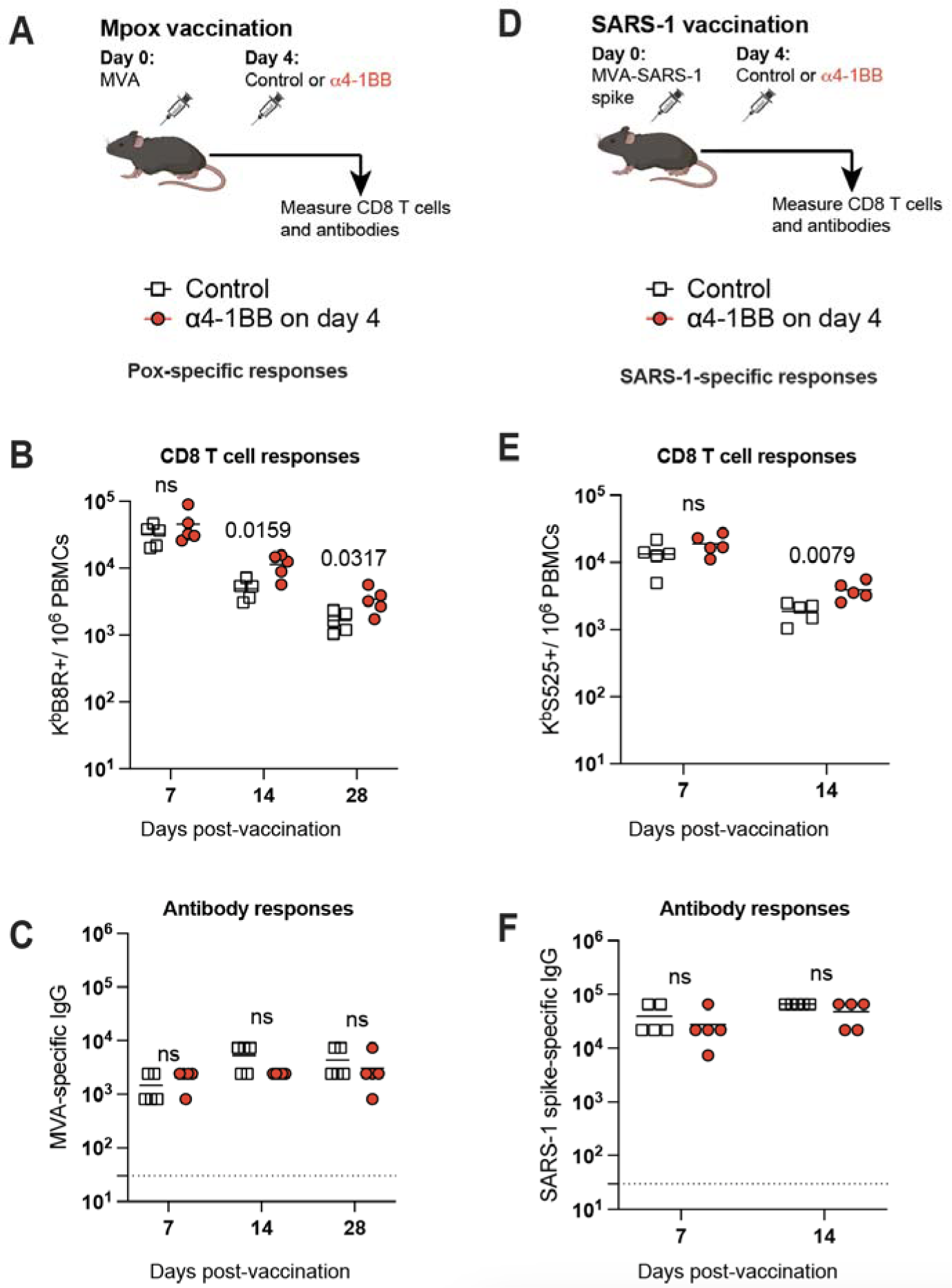
Generalizability to another vaccine platform: a poxvirus vector. **(A)** Experimental outline for evaluating whether treatment with α4-1BB improves immune responses elicited by an MVA vaccine. **(B)** Summary of MVA-specific CD8 T cell responses in PBMCs. **(C)** Summary of MVA-specific antibody responses in sera. **(D)** Experimental outline for evaluating whether treatment with α4-1BB improves immune responses elicited by an MVA-SARS-CoV-1 vaccine. **(E)** Summary of SARS-CoV-1 specific CD8 T cell responses in PBMCs. **(F)** Summary of SARS-CoV-1-specific antibody responses in sera. Mice were vaccinated with 10^7^ PFU of the respective MVA vector followed by treatment with 50 μg of α4-1BB or control antibodies at day 4. Data are from one experiment, n=5 per group. Indicated *P* values were determined by parametric test (unpaired t test).

### Kinetics of 4-1BB expression

The data above suggest that extending the time interval between antigen recognition and costimulation may be beneficial for CD8 T cell responses. In other words, triggering 4-1BB costimulation on the same day of vaccination did not improve CD8 T cells, whereas triggering 4-1BB costimulation at day 4 of vaccination did improve CD8 T cells. To better understand this time-dependent effect, we measured the expression of 4-1BB on antigen-specific CD8 T cells. During the acute phase of the immune response, the precursor frequency of virus-specific CD8 T cells is very low, rendering it difficult to quantify 4-1BB levels on antigen-specific CD8 T cells. Thus, we utilized a chimera model in which a high number of virus-specific CD8 T cells are transferred into recipient mice, allowing us to measure 4-1BB kinetics at early timepoints post-vaccination. In brief, we adoptively transferred Thy1.1+ congenically labeled CD8 T cells specific for LCMV (P14 cells) into naïve recipient mice via the intravenous route. One day after, the recipient mice were vaccinated intramuscularly with the mRNA-GP vaccine and 4-1BB expression was measured on donor P14 cells (**Fig. S4A**). 4-1BB expression on virus-specific CD8 T cells was high at day 4 post-vaccination and returned to baseline levels by day 7 of vaccination (**Fig. S4B**). Similar “zig-zag” expression kinetics were observed with other costimulatory molecules, including CD27, CD28, and OX40, which showed highest expression at day 4 post-vaccination (**Fig. S4B**).

Finally, we examined the effects of triggering 4-1BB costimulation at later timepoints post-vaccination (day 14, which corresponds to the contraction phase of the immune response). Treatment of mice with 4-1BB costimulatory antibodies at day 14 post-vaccination did not elicit any discernable effects on CD8 T cells or antibody responses (**Fig. S5A-S5C**). These findings demonstrate that 4-1BB costimulation has time-dependent effects on vaccine responses, and that modulation of the 4-1BB pathway, specifically during the early effector phase, could serve as a potent adjuvant to improve vaccine efficacy.

## DISCUSSION

mRNA vaccines have been administered to millions of people worldwide and have shown high efficacy at preventing severe disease and death caused by SARS-CoV-2 infection. However, these vaccines do not confer sterilizing immunity, and require multiple booster shots. This has motivated the efforts to develop improved mRNA vaccine regimens. In this study, we interrogated whether mRNA vaccines could be improved by triggering 4-1BB, a costimulatory molecule involved in T cell activation. 4-1BB costimulatory antibodies have been clinically tested in autoimmunity and cancer immunotherapy ^17, 18^, and the signaling molecules involved in 4-1BB costimulation are included in chimeric antigen receptor (CAR) T cell therapies ^29^. Although 4-1BB costimulation is known to play an important role in immune activation, it could also cause immunosuppression in certain contexts, and it is unclear how the timing of 4-1BB costimulation determines these opposite outcomes. A prior study reported that when 4-1BB costimulatory antibodies are administered at day 1 of LCMV infection, both T cell and antibody responses are impaired ^20^. Similarly, other studies have demonstrated impaired antibody responses associated with 4-1BB costimulatory antibodies, when these are administered concurrently with vaccination ^30, 31^. Consistent with these prior studies, we also demonstrate that administration of 4-1BB costimulatory antibodies on the same day of vaccination impairs antibody responses. These immunosuppressive effects may not be exclusive to 4-1BB. A prior study demonstrated that after LCMV infection, administering CD40 costimulatory antibodies leads to impaired immune responses ^32^. Collectively, these results suggest that concurrent provision of Signal 1 and Signal 2 may not be optimal for immune responses, and that temporally separating these signals may be necessary to fully unleash the immunostimulatory effects of costimulation.

Antigen recognition is metaphorically analog to inserting the key to turn on a car, while costimulation is analog to stepping on the accelerator. Employing this classical analogy, inserting the key and stepping on the accelerator at the same time can lead to “flooding of the engine.” This concept led us to hypothesize that extending the time interval between antigen recognition and costimulation would allow CD8 T cells to “warm up” and upregulate their costimulatory receptors, rendering them more responsive to subsequent costimulation. Our kinetics data demonstrates the inducible nature of 4-1BB and other costimulatory molecules following mRNA vaccination, and are consistent with prior studies using viral infection models, showing that costimulatory molecules like 4-1BB are transiently induced by TCR signaling ^33^.

In the current study, we make the novel observation that a temporal separation between immunization and costimulation improves vaccine efficacy. This time-dependent effect could be explained by the fact that 4-1BB is an “inducible” costimulatory receptor. 4-1BB costimulatory antibodies may not be able to engage many 4-1BB receptors on naïve CD8 T cells since these cells have not yet upregulated 4-1BB on their surface. However, treatment with 4-1BB costimulatory antibodies at day 4 (when CD8 T cells express high levels of 4-1BB) may trigger more potent costimulatory signaling, leading to more robust expansion of CD8 T cells. The inducible antigen-dependent nature of 4-1BB as well as other costimulatory receptors likely ensures the proper sequence of signaling events, precluding “out-of-order” signaling (e.g. costimulation preceding antigen recognition).

We also examined the specific CD8 T cell subsets that were affected by 4-1BB costimulation. We show that 4-1BB costimulation at day 4 resulted in a significant increase in effector CD8 T cells, as well as effector memory CD8 T cells, while exerting no significant effects on central memory CD8 T cells. These data are consistent with a prior study from Watts et al. that suggests that 4-1BB is important for the persistence of effector CD8 T cells in tissues ^34^. A cardinal feature of effector and effector memory CD8 T cells is their “response-ready” state ^26^ which provides rapid sterilizing protection upon breakthrough infection, but these cells have a shorter lifespan relative to central memory CD8 T cells. Notwithstanding, we detected increased CD8 T cell responses up to 70 days post-vaccination in mice that received 4-1BB costimulatory antibodies, suggesting long-term enhancement of CD8 T cell responses by 4-1BB costimulation. Future studies will examine the durability of immune responses over longer periods of time.

Our data with multiple challenge models show that 4-1BB costimulation at day 4 improves vaccine-mediated protection against breakthrough infection, but the level of immune protection varied depending on the challenge model. For example, in the LCMV and listeria challenge models, most animals that received 4-1BB after vaccination exhibited complete sterilizing immunity upon challenge, with no evidence of infection, whereas in the SARS-CoV-2 challenge model the improvement was more modest relative to control vaccination. These differences in protection across various challenge models may be attributed to the relative roles of CD8 T cells versus antibodies in clearing specific infections. In the case of intracellular pathogens like arenaviruses or listeria, CD8 T cells are indispensable for pathogen clearance, and given that 4-1BB costimulation enhances CD8 T cells, this likely explains the more drastic immunologic effects observed with LCMV and listeria.

A potential limitation of our findings is the need for vaccinees to undergo an additional injection with costimulatory antibodies, 4 days after initial vaccination. To address this logistical challenge, future studies should examine the effects of encapsulating 4-1BB costimulatory antibodies in slow-release formulations, which could be co-administered with the mRNA vaccine on the same day. Another consideration is that 4-1BB costimulatory antibodies can elicit severe inflammation, which is a reason why 4-1BB costimulatory antibodies have not yet been licensed ^35^. However, in our studies we used a single low dose of 4-1BB costimulatory antibodies, and we showed that the effects on immune responses were similar to when we administered high, repetitive doses, suggesting that a small single dose only at day 4 post-vaccination is sufficient to induce a durable improvement in vaccine immunity.

The classical 2-signal model established by Bretscher et al. postulates that T cell activation is dependent on two consecutive signals (antigen recognition and costimulation) also known as Signal 1 and Signal 2, respectively. For decades, this model has been a blueprint to understand how T cell responses are generated and has had broad implications for the development of immunotherapies and vaccine adjuvants, aimed at enhancing costimulation. However, it is currently unknown whether there is an ideal timing gap between these two signals. Our study brings more clarity to this issue, as we show the benefits of temporally separating antigen recognition and 4-1BB costimulation by several days. Future studies will examine the effects of delayed costimulation with other costimulatory pathways, such as CD28, CD27 and OX40, which also showed high expression at day 4 post-vaccination. Prior studies have also examined the effects of triggering costimulatory pathways in cancer immunotherapies and vaccines, but clinical efficacy has been limited ^36^. A prior study that focused on patients receiving CD27 costimulatory antibodies for cancer treatment revealed a notable outcome: the patient with highest CD27 expression achieved complete response after treatment with CD27 costimulatory antibodies ^37^. This finding suggests that the efficacy of costimulatory antibodies can be dictated by the expression levels of costimulatory receptors. In summary, our results highlight a critical time-dependent effect of the 4-1BB pathway on vaccine immunity, warranting the clinical evaluation of 4-1BB agonists to improve vaccine efficacy.

## ACKNOWLEDGEMENTS

We thank the late Dr. Robert Mittler for discussions on how 4-1BB regulates immune responses. We also thank Dr. Gordon Freeman for discussions and Dr. Thomas Gallagher for help designing the mRNA vaccine against the common cold coronavirus, OC43. Dr. This work was possible with a grant from the National Institute on Drug Abuse (NIDA, DP2DA051912) to P.P.M.

## AUTHOR CONTRIBUTIONS

P.P.M. and S.S. designed the experiments. S.S. performed most of the experiments. T.D., N.I. and B.A. helped with some of the immunogenicity experiments. S.F. analyzed the gene expression data. J.R. performed the SARS-CoV-2 challenges. P.P.M. and S.S. wrote the manuscript with feedback from all authors.

## Figure Legends

**Supp. Fig. 1.**
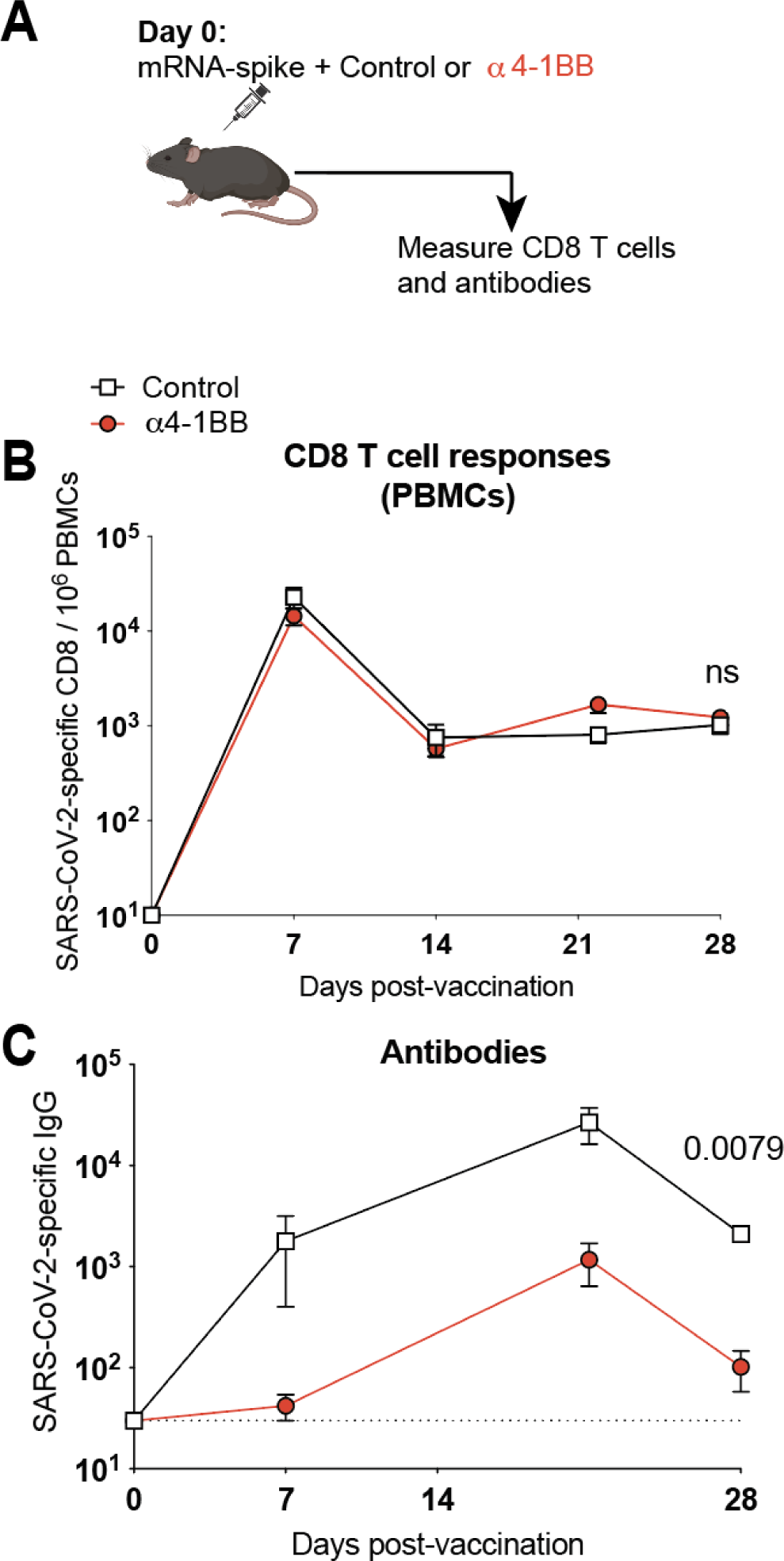
4-1BB costimulation during the acute priming phase does not result in improvement of immune responses following mRNA-SARS-CoV-2 vaccination. **(A)** Experimental outline for evaluating whether treatment with α4-1BB improves immune responses elicited by an mRNA-SARS-CoV-2 vaccine in C57BL/6 mice. Mice were immunized with 3 μg of a SARS-CoV-2 mRNA spike vaccine followed by α4-1BB or control antibodies at day 0 (on the same of vaccination). (B) Summary of SARS-CoV-2-specific CD8 T cell responses in PBMCs. (C) Summary of SARS-CoV-2-specific antibody responses in sera. Data are from one experiment, n=5 per group. Indicated *P* values were determined by parametric test (unpaired t test).

**Supp Fig. 2.**
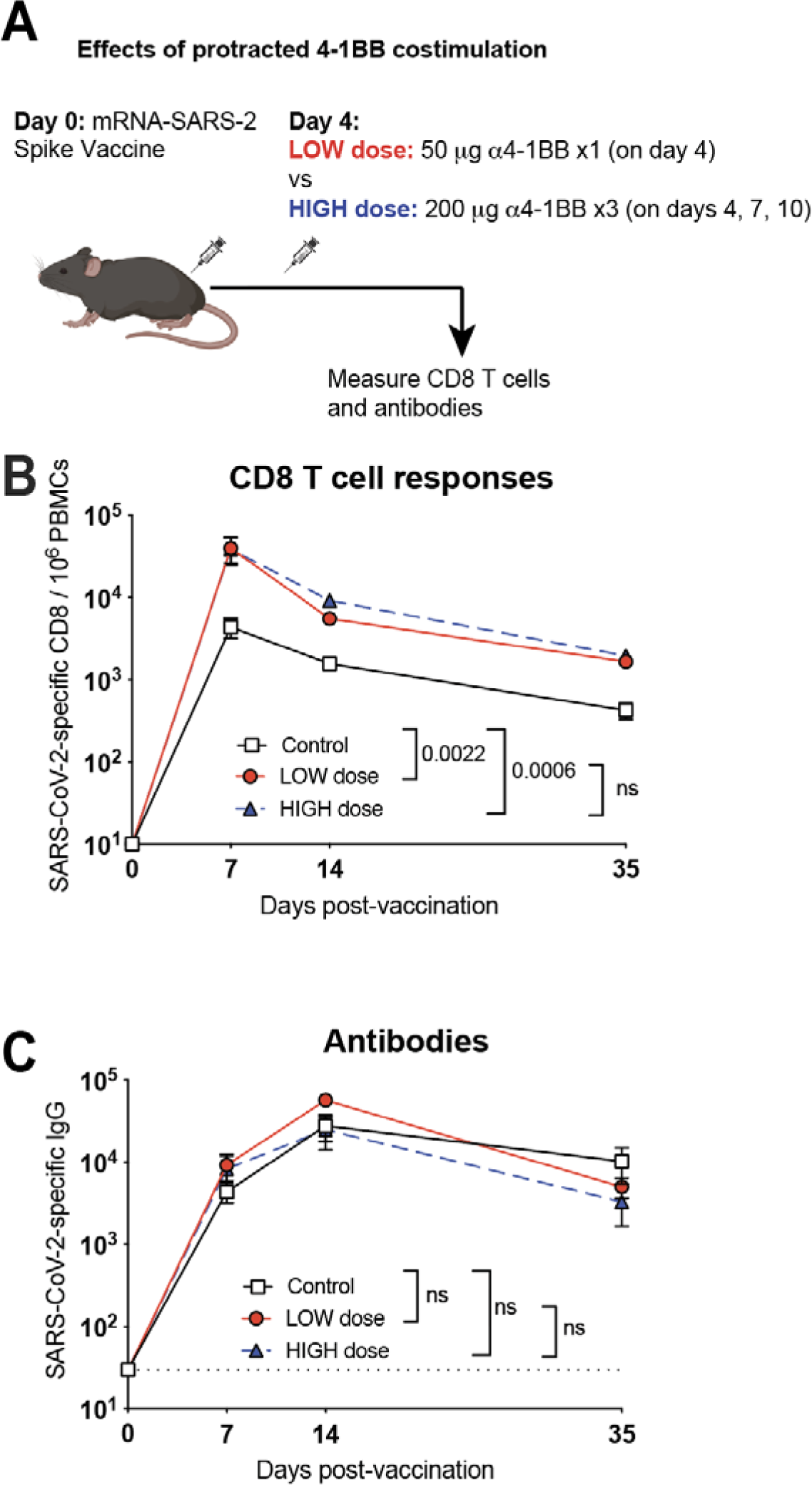
Long-term 4-1BB costimulation does not result in superior immune responses relative to short-term 4-1BB costimulation. **(A)** Experimental outline for evaluating the effect of a high dose α4-1BB treatment. Mice were vaccinated with 3 μg of an mRNA-SARS-CoV-2 vaccine. At day 4, one group of mice received a single dose of 50 μg of α4-1BB (low dose); and another group of mice received 200 μg of α4-1BB multiple times (high dose). (**B**) Summary of SARS-CoV-2-specific CD8 T cell responses in PBMCs. (**C**) Summary of SARS-CoV-2-specific antibody responses in sera. Data are from one experiment with n=5 per group. Indicated *P* values were determined by 2-way ANOVA (Dunnett’s multiple comparisons tests) at the last time point.

**Supp Fig. 3.**
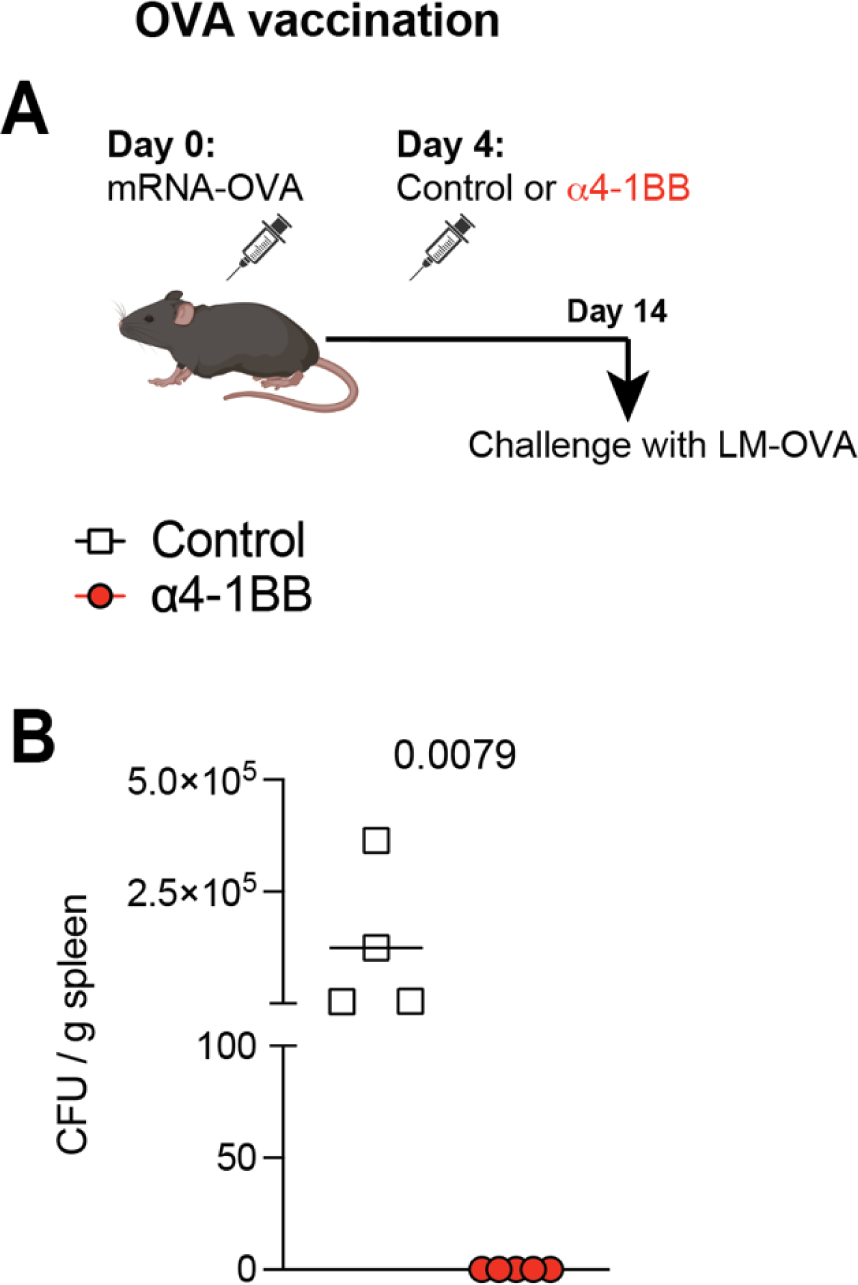
4-1BB costimulation following immunization with an mRNA-based listeria vaccine confers sterilizing immunity. **(A)** Experimental outline for evaluating whether treatment with α4-1BB improves immune responses elicited by an mRNA-OVA vaccine against a listeria-OVA (LM-OVA) challenge. Mice were vaccinated with 3 μg of an mRNA-OVA vaccine followed by treatment with 50 μg of α4-1BB or control antibodies at day 4. At day 14, they were challenged intravenously with a supra-lethal dose of LM-OVA (10^7^ CFU) and bacterial loads were quantified. (**B**) Summary of bacterial loads in spleen at day 3 post-challenge. Data are from one experiment, n=4-5 per group. Indicated *P* values were determined by parametric test (unpaired t test).

**Supp. Fig. 4.**
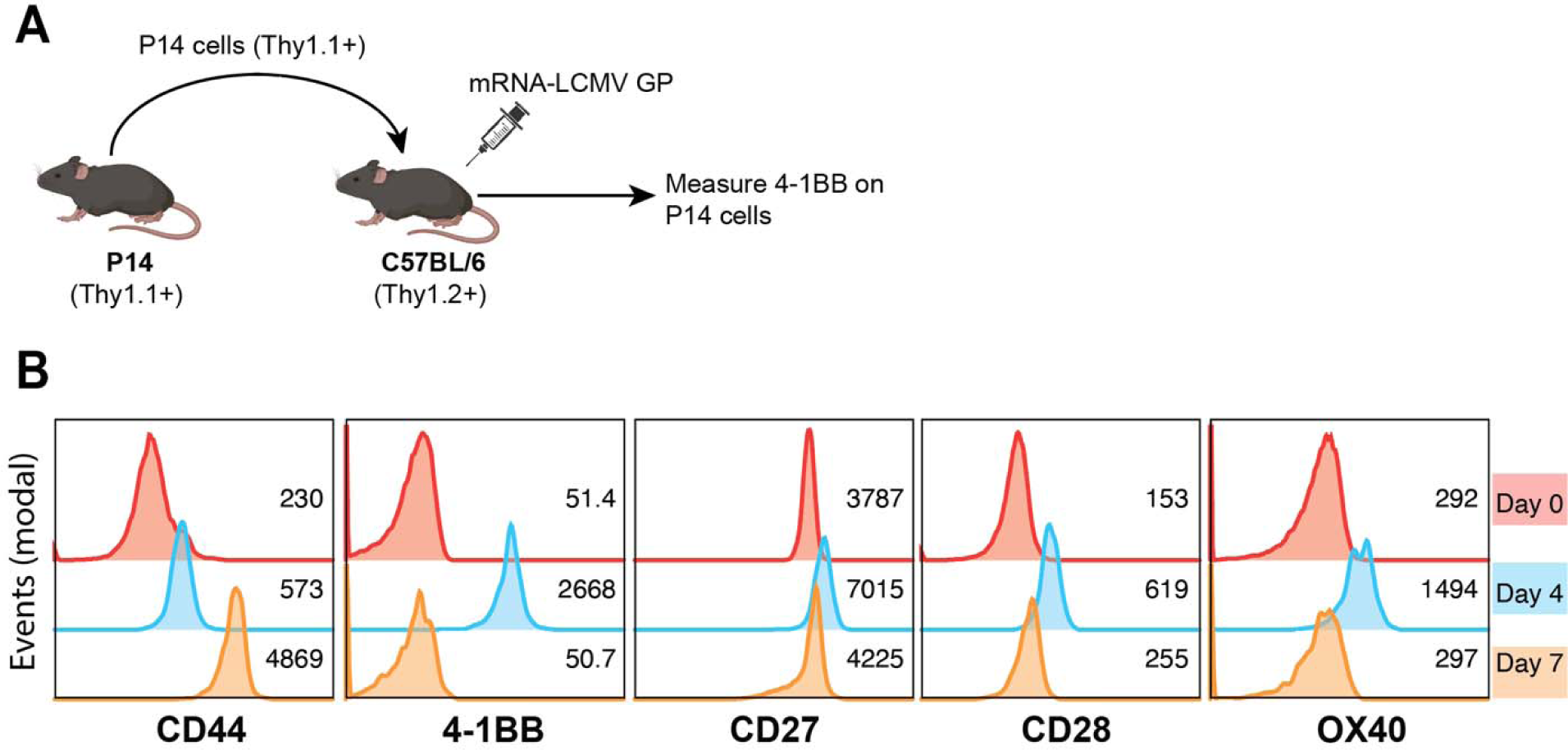
4-1BB is induced on virus-specific CD8 T cells after mRNA vaccination. **(A)** Experimental outline for evaluating 4-1BB expression following mRNA vaccination. ∼35,000 congenically-labeled (Thy1.1+) CD8 T cells from a P14 transgenic mouse were transferred intravenously into a Thy1.2+ C57BL/6 mouse. One day after P14 transfer, C57BL/6 mice were immunized with 3 μg of a mRNA-LCMV GP vaccine, and 4-1BB was measured on virus-specific CD8 T cells at various time points. **(B)** Representative histograms showing 4-1BB and other costimulatory molecule expression on virus-specific (P14) CD8 T cells. CD44 is shown to visualize activationWe utilized this P14 chimera model using high number of P14 cells to allow us to detect 4-1BB expression on virus-specific CD8 T cells at hyperacute points; note that endogenous virus-specific CD8 T cells cannot be detected at hyperacute time points due to their low precursor frequency. Experiment was performed 2 times with n=3 per group, showing similar results (peak of 4-1BB expression at day 4 post-vaccination).

**Supp. Fig. 5.**
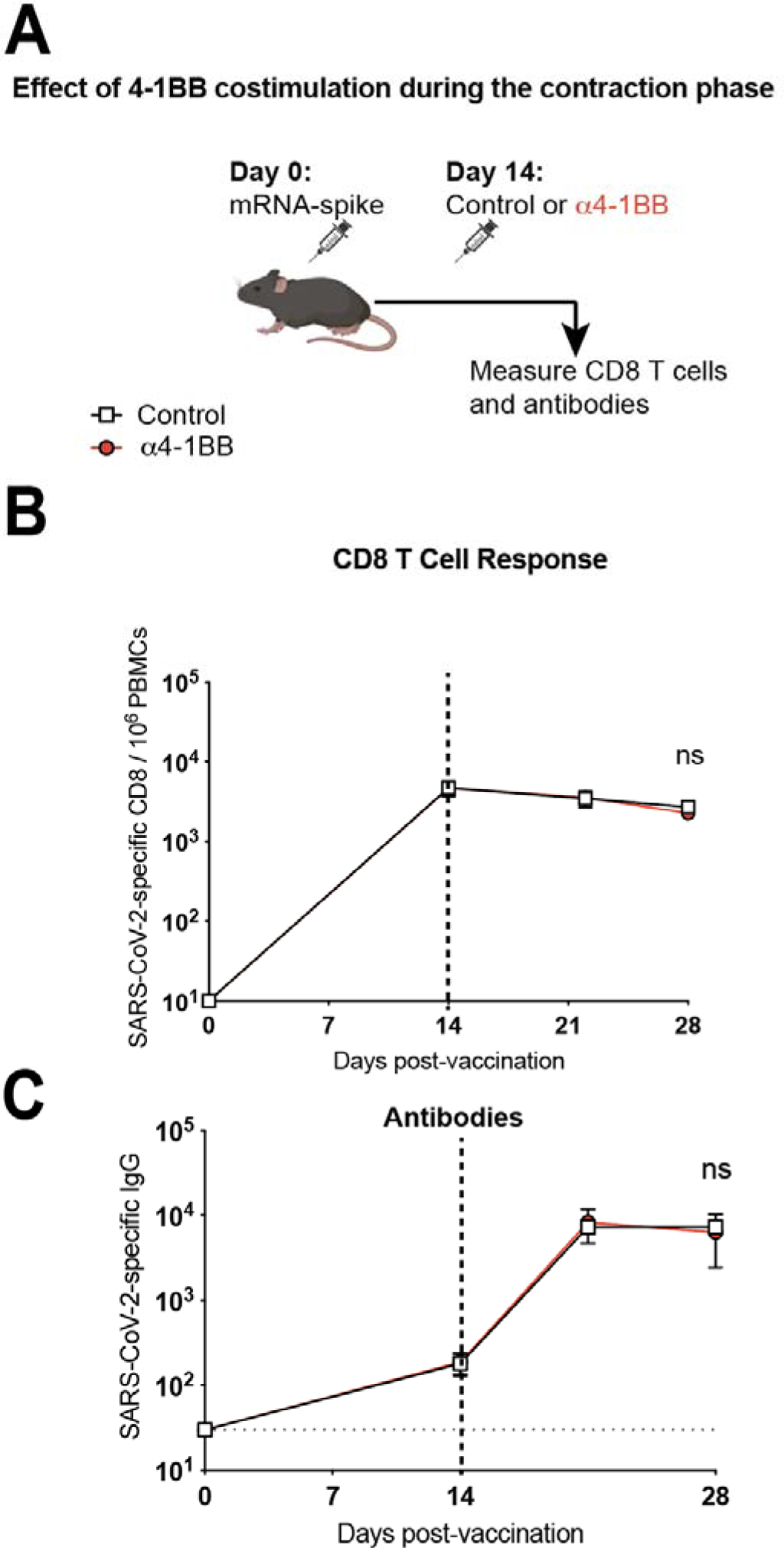
4-1BB costimulation during the contraction phase does not result in improvement of immune responses following mRNA-SARS-CoV-2 vaccination. **(A)** Experimental outline for evaluating whether treatment with α4-1BB improves immune responses elicited by an mRNA-SARS-CoV-2 vaccine in C57BL/6 mice. Mice were immunized with 3 μg of a SARS-CoV-2 mRNA spike vaccine followed by treatment with α4-1BB or control antibodies at day 14. **(B)** Summary of SARS-CoV-2-specific CD8 T cell responses in PBMCs. (**C**) Summary of SARS-CoV-2-specific antibody responses in sera. Data are from one experiment, n=5 per group. Indicated *P* values were determined by parametric test (unpaired t test).

## MATERIALS AND METHODS

### Mice and vaccinations

6-8-week-old C57BL/6 mice were used. Mice were purchased from Jackson laboratories (approximately half males and half females). Mice were immunized intramuscularly with mRNA-LNPs (made in house) or MVA vectors (from Dr. Bernard Moss, NIH) diluted in sterile PBS. Mice received α4-1BB agonistic antibody (clone 3H3, BioXcell) or IgG control antibody (clone 2A3, BioXcell) intraperitoneal (50 μg or 200 μg on the indicated days) diluted in sterile PBS. Mice were housed at Northwestern University’s Center for Comparative Medicine (CCM) or University of Illinois at Chicago (UIC). All mouse experiments were performed with approval of the Northwestern University Institutional Animal Care and Use Committee (IACUC).

### Reagents, flow cytometry and equipment

Single cell suspensions were obtained from PBMCs, and various tissues as described previously. Dead cells were gated out using Live/Dead fixable dead cell stain (Invitrogen). SARS-CoV-2 spike and SF162 peptide pools used for intracellular cytokine staining (ICS) and these were obtained from BEI Resources. Biotinylated MHC class I monomers (K^b^VL8, sequence VNFNFNGL; D^b^GP33, sequence KAVYNFATC; D^b^GP276, sequence SGVENPGGYCL; OVA, sequence SIINFEKL; K^b^B8R, sequence TSYKFESV) were used for detecting virus-specific CD8 T cells, and were obtained from the NIH tetramer facility at Emory University. Cells were stained with fluorescently-labeled antibodies against CD8α (53-6.7 on PerCP-Cy5.5), CD44 (IM7 on FITC), CD62L (MEL-14 on PE-Cy7), CD127 (A7R34 on Pacific Blue), and tetramers (APC). TNFα (MP6-XT22 on PE-Cy7), IFNγ (XMG1.2 on APC), or Ki67 (SolA15 on PE-Cy7). Fluorescently-labeled antibodies were purchased from BD Pharmingen, except for anti-CD127 and anti-CD44 (which were from Biolegend). Flow cytometry samples were acquired with a Becton Dickinson Canto II or an LSRII and analyzed using FlowJo v10 (Treestar).

### SARS-CoV-2 spike, SARS-CoV-1 spike, OVA, HIV (SF162) envelope, and MVA lysate-specific ELISA

Binding antibody titers were measured using ELISA as described previously ^4, 23–25, 38–40^. In brief, 96-well flat bottom plates MaxiSorp (Thermo Scientific) were coated with 0.1μ/well of the respective spike protein, for 48 hr at 4°C. For detection of MVA-specific antibody responses, MVA lysates were used as coating antigen (incubated for 48 hr at room temperature). Plates were washed with PBS + 0.05% Tween-20. Blocking was performed for 4 hr at room temperature with 200 μL of PBS + 0.05% Tween-20 + bovine serum albumin. 6μL of sera were added to 144 μL of blocking solution in first column of plate, 1:3 serial dilutions were performed until row 12 for each sample and plates were incubated for 60 minutes at room temperature. Plates were washed three times followed by addition of goat anti-mouse IgG horseradish peroxidase conjugated (Southern Biotech) diluted in blocking solution (1:5000), at 100 μL/well and incubated for 60 minutes at room temperature. Plates were washed three times and 100 μL /well of Sure Blue substrate (Sera Care) was added for approximately 8 minutes. Reaction was stopped using 100 μL/well of KPL TMB stop solution (Sera Care). Absorbance was measured at 450 nm using a Spectramax Plus 384 (Molecular Devices). SARS-CoV-2 spike protein was produced in-house using a plasmid produced under HHSN272201400008C and obtained from BEI Resources, NIAID, NIH: vector pCAGGS containing the SARS-related coronavirus 2; Wuhan-Hu-1 spike glycoprotein gene (soluble, stabilized); NR-52394. SARS-CoV-1 spike protein was obtained through BEI Resources, NIAID, NIH: SARS-CoV Spike (S) Protein deltaTM, Recombinant from Baculovirus, NR-722. HIV-SF162 protein was obtained through the NIH AIDS Reagent Program, Division of AIDS, NIAID, NIH: Human Immunodeficiency Virus Type 1 SF162 gp140 Trimer Protein, Recombinant from HEK293T Cells, ARP-12026, contributed by Dr. Leo Stamatatos. **OVA protein was produced in-house using a plasmid produced under (details from TD to follow).**

### mRNA-LNP vaccines

We synthesized mRNA vaccines encoding for the codon-optimized SARS-CoV-2 spike protein from USA-WA1/2020, OC43 spike protein, OVA from the SERPINB14 gene, HIV-1 SF162 envelope protein, or the LCMV GP. Constructs were purchased from Integrated DNA Technologies (IDT) or Genscript, and contained a T7 promoter site for *in vitro* transcription of mRNA. The sequences of the 5′- and -3′′-UTRs were identical to those used in a previous publication.^44^ All mRNAs were encapsulated into lipid nanoparticles using the NanoAssemblr Benchtop system (Precision NanoSystems) and confirmed to have similar encapsulation efficiency (∼95%). mRNA was diluted in Formulation Buffer (Catalog # NWW0043, Precision NanoSystems) to 0.17 mg/mL and then run through a laminar flow cartridge with GenVoy ILM encapsulation lipids (Catalog # NWW0041, Precision NanoSystems) with N/P (Lipid mix/mRNA ratio of 4) at a flow ratio of 3:1 (RNA: GenVoy-ILM), with a total flow rate of 12 mL/min, to produce mRNA– lipid nanoparticles (mRNA-LNPs). mRNA-LNPs were evaluated for encapsulation efficiency and mRNA concentration using RiboGreen assay using the Quant-iT RiboGreen RNA Assay Kit (Catalog # <u>R11490</u>, Invitrogen, Thermo Fisher Scientific).

### RNA-Seq Data Acquisition and Analysis

C57BL/6 mice were immunized with a 3 μg of mRNA-SARS-CoV-2 spike, and at day 4, treated with 4-1BB agonistic antibody. At day 7, splenic CD8 T cells were MACS-sorted with a MACS negative selection kit (STEMCELL). Purified CD8 T cells were stained with K^b^ VL8 tetramer, live dead stain, and antibodies for CD8 and CD44 to gate on virus-specific CD8 T cells. Live, CD8+, CD44+, K^b^VL8+ cells were FACS-sorted to ∼99% purity on a FACS Aria cytometer (BD Biosciences) and delivered to Admera Health Biopharma for RNA extraction using Illumina 2×150 and RNA-seq using SMARTseq V4 with NexteraXT kit. After the library was sequenced, the output file in BCL format was converted to FASTQ files and aligned to mouse genome to generate a matrix file using the Cell Ranger pipeline (10X Genomics). These upstream QC steps were performed by Dr. Slim Fourati at Northwestern University. Further analyses were performed in R using the Seurat package v4.0, as previously described ^41^. Terminal effector gene signatures were derived using the edgeR package ^42^, comparing effector memory to terminal effector CD8 T cells ^43^. Clusters representing less than 4% of each population were excluded from downstream analyses. RNA-Seq accession data were deposited in the NCBI’s Gene Expression Omnibus (GEO) database; provided after peer review).

### Adoptive transfer of P14 cells

CD8+ T cells from Thy1.1^+^ P14 mice (PBMCs) were enriched using a CD8 MACS-negative selection kit (STEMCELL Technologies). ∼35,000 P14 CD8^+^ T cells were transferred intravenously into naïve Thy1.2^+^ C57BL/6 recipient mice. Recipient mice were vaccinated intramuscularly with 3 μg of the mRNA-LCMV vaccine 24 hours later. PBMCs were collected at various time points to measure 4-1BB expression on donor (Thy1.1+) P14 T cells by flow cytometry.

### SARS-CoV-2 virus and other challenge models

SARS-Related Coronavirus 2, Isolate USA-WA1/2020, NR-52281 was deposited by the Centers for Disease Control and Prevention and obtained through BEI Resources, NIAID, NIH. This virus was propagated and tittered on Vero-E6 cells (ATCC). Vero-E6 cells were passaged in DMEM with 10% Fetal bovine serum (FBS) and Glutamax. Cells less than 20 passages were used for all studies. Virus stocks were expanded in Vero-E6 cells following a low MOI (0.01) inoculation and harvested after 96 hr. Titers were determined by plaque assay on Vero-E6 cell monolayers. Viral stocks were used after a single expansion (passage = 1) to prevent genetic drift. K18-hACE2 mice were anesthetized with isoflurane and challenged with 5 × 10^4^ PFU of SARS-CoV-2 intranasally. Mouse challenges were performed at the University of Illinois at Chicago (UIC) following BL3 guidelines with approval by the UIC Institutional Animal Care and Use Committee (IACUC). LCMV Cl-13 challenges were intravenously at 2 x 10^6^ PFU, and listeria (LM-OVA) challenges were intravenously at 10^7^ CFU.

### SARS-CoV-2 quantification

Lungs were collected from infected mice and RNA was isolated with the Zymo 96-well RNA isolation kit (Catalog #: R1052) following the manufacturer’s protocol. SARS-CoV-2 viral burden was measured by RT-qPCR using Taqman primer and probe sets from IDT with the following sequences: Forward 5′ GACCCCAAAATCAGCGAAAT-3′, Reverse 5′ TCTGGTTACTGCCAGTTGAATCTG-3′, Probe 5′ ACCCCGCATTACGTTTGGTGGACC-3′. A SARS-CoV-2 copy number control was obtained from BEI (NR-52358) and used to quantify SARS-CoV-2 genomes.

### LCMV virus quantification

Seed stock of LCMV Cl-13 was obtained from Dr. Rafi Ahmed’s laboratory. The virus was propagated and tittered on BHK21 cells (ATCC). BHK21 cells were passaged in DMEM with 10% Fetal bovine serum (FBS). Cells were inoculated with a low MOI (0.1) in 1% DMEM and incubated for 72 hr. Titers were determined by plaque assay on Vero-E6 cell monolayers. Sera and spleen were collected at various time points post-challenge. Infectious viral titers were determined by plaque assay using Vero-E6 cells. 5 x 10^5^ Vero-E6 cells per well were seeded in 6-well plates in 10% DMEM media, monolayer was 90-100% confluent after 24 hr. Spleen was homogenized using a standard TissueRuptur homogenizer, and 10-fold serial dilution of tissue were made then transferred dropwise onto cell monolayer. Sera dilutions were created in 10% DMEM media and added dropwise on cell monolayer. 6-well plates were placed in 37°C 5% CO_2_ incubator for 1 hr and manually rocked every 10 minutes. A 1:1 agarose/2×199 monolayer was dispensed after 1 hr incubation and plates were incubated at 37°C 5% CO_2_ for 96 hr. After 96 hr a 1:50 1% neutral red solution was added to a 1:1 agarose/2×199 mixture and overlayed onto plates. Plaques were counted the following day after agar overlay removal.

### Listeria Quantification

Spleens were collected from infected mice at day 3 post-challenge. Bacterial titers were quantified by homogenizing tissues through a 42um strainer and resuspended in 1% Triton. 10-fold serial dilutions were created in 1% Triton and added dropwise onto 6-well BHK agar plates. Plates were manually rocked then incubated at 37°C 5% CO2 for 24 hours. Colonies were counted the next day.

### Statistical analysis

Statistical tests used are indicated on each figure legend. Dashed lines in data figures represent limit of detection. Data were analyzed using Prism version 10 (Graphpad).

### Competing Interests

The authors declare no competing interests.

## Notes

### Competing Interest Statement

The authors have declared no competing interest.

